# Effects of *Nf1* on sleep behavior are mediated through starvation caused by deficits in SARM1 dependent NAD+ metabolism

**DOI:** 10.1101/2024.09.14.612058

**Authors:** Folasade A. Sofela, Mariela Lopez Valencia, Thomas A. Jongens, Amita Sehgal

## Abstract

Neurofibromatosis 1 (NF1) is a relatively common autosomal dominant disease which predisposes to the formation of tumors, and is also associated with behavioral phenotypes, including sleep disturbances. As loss of the NF1 protein has been recently associated with metabolic dysfunction, we explored the relationship between metabolic and behavioral phenotypes through metabolomic analysis of *Drosophila Nf1*-null mutants. *Nf1*-null mutants exhibit a metabolic signature indicative of starvation, with diminished metabolites related to glucose, glycogen, and fatty acid processing and increased mRNA of *Akh*, a hormone that promotes foraging during starvation. Reduced sleep in *Nf1*-null mutants was rescued by genetic manipulation of the AKH pathway and by a high-sucrose diet, which also partially corrected hypolipidemia, suggesting that sleep loss is due to starvation-induced foraging. Interestingly, behavioral phenotypes can be recapitulated by loss of NF1 only in the periphery and trace to mitochondrial defects that include elevated levels of the NADase SARM1. Indeed, inhibition of SARM1 activity rescues sleep behavior in *Nf1*-null flies. These findings suggest a novel connection between loss of NF1 and mitochondrial dysfunction caused by SARM1 hyperactivation, setting the scene for new pharmacological and dietary approaches that could provide relief to NF1 patients.

## Introduction

Neurofibromatosis type 1 (NF1) is a genetic disorder associated with several physiological abnormalities across multiple systems, including cognitive impairment, musculoskeletal abnormalities, and increased susceptibility to benign and malignant nervous system tumors (*1*). NF1 affects 1:2000 to 1:3500 people, and individuals with NF1 typically develop persistent clinical problems that accumulate throughout their shortened lifespan (*2*, *3*). Sleep disturbances are a common symptom in NF1, with up to 70% of children with the disease experiencing sleep problems, especially difficulty initiating and maintaining sleep (*4–7*). Attention Deficit Hyperactivity Disorder (ADHD) is also a common comorbidity in NF1, with over 50% of children with NF1 meeting the CPRSLR diagnostic criteria for ADHD symptomatology, an incidence nearly ten times that of the general population. (*8*).

NF1 is caused by inactivating mutations in *Nf1*, the gene that encodes for neurofibromin on chromosome 17 (*9*). Neurofibromin is a Ras GTPase activating protein whose known function is to inactivate the Ras oncogene through GTP hydrolysis (*10*, *11*).

Thus, the consequence of inactivating mutations in *Nf1* is unregulated Ras-MAPK signaling leading to tumor formation. However, the mechanisms underlying behavioral abnormalities in NF1 are still largely unclear. Notably, metabolic dysfunction, especially defects in energy metabolism, glucose handling and insulin signaling, has recently emerged as a characteristic of NF1, but it has been largely understudied as a possible cause or exacerbating factor in NF1 symptomology (*12–15*).

*Drosophila* models of NF1 phenocopy many aspects of the human disorder, including sleep disturbance (*16*) and increased metabolic rate (*17*). Thus, these models have been invaluable in the study of the disease, its associated pathologies, and mechanisms underlying the disease. Here, we aimed to shed light on the relationship between NF1, sleep, and metabolism using *Drosophila* models of NF1. Through a series of experiments, we show that loss of *Nf1* results in a severe energy deficit that leads to hyperactivity and sleep loss secondary to organismal starvation. Additionally, our results indicate that the source of this energy crisis is a defect in mitochondrial NAD+ and Ca2+ metabolism fueled by the hyperactivation of the mitochondrial NAD+ hydrolase SARM1. These results have important implications for the development of new treatments for NF1 and may lead to a deeper understanding of the disease.

### Metabolomic Study of *Nf1*-null *Drosophila*

We conducted a metabolomic study of two genotypically distinct *Drosophila Nf1*-null mutant alleles (P1 and P2), to determine the metabolic consequences of loss of NF1. A principal component analysis of the metabolomic data revealed clear, distinct differences between mutants and age matched controls, indicating that there is indeed a specific and significant metabolic phenotype that can be attributed to loss of NF1 (Fig. 1a). Previous studies showed that in human and mouse, loss of *Nf1* increases glucose tolerance and insulin sensitivity while lowering fasting blood glucose (*12–15*). Consistent with these findings, glucose and lipid species were dramatically reduced in both *Nf1* mutants, as compared to their isogenic age-matched control *K33* (Fig. 1b). Interestingly, levels of glycolytic end products like lactate, pyruvate, and alanine as well as glycogen intermediates like maltotriose and maltotetraose were also significantly reduced in both *Nf1* mutants, suggesting that the metabolic phenotype of *Nf1* represents a generalized defect in energy metabolism (Fig.1b) b, Extended Data Fig.1 b-g). In addition, a whole-body reduction in the levels of triglycerides and ATP suggests that the energy deficit is not simply a result of reduced glucose levels, but rather a general defect in energy production (Fig.2a, b). Importantly, these results reveal a role for NF1 in regulating energy metabolism that may contribute to the smaller size of NF1 patients and mutant flies.

**Figure 1.**
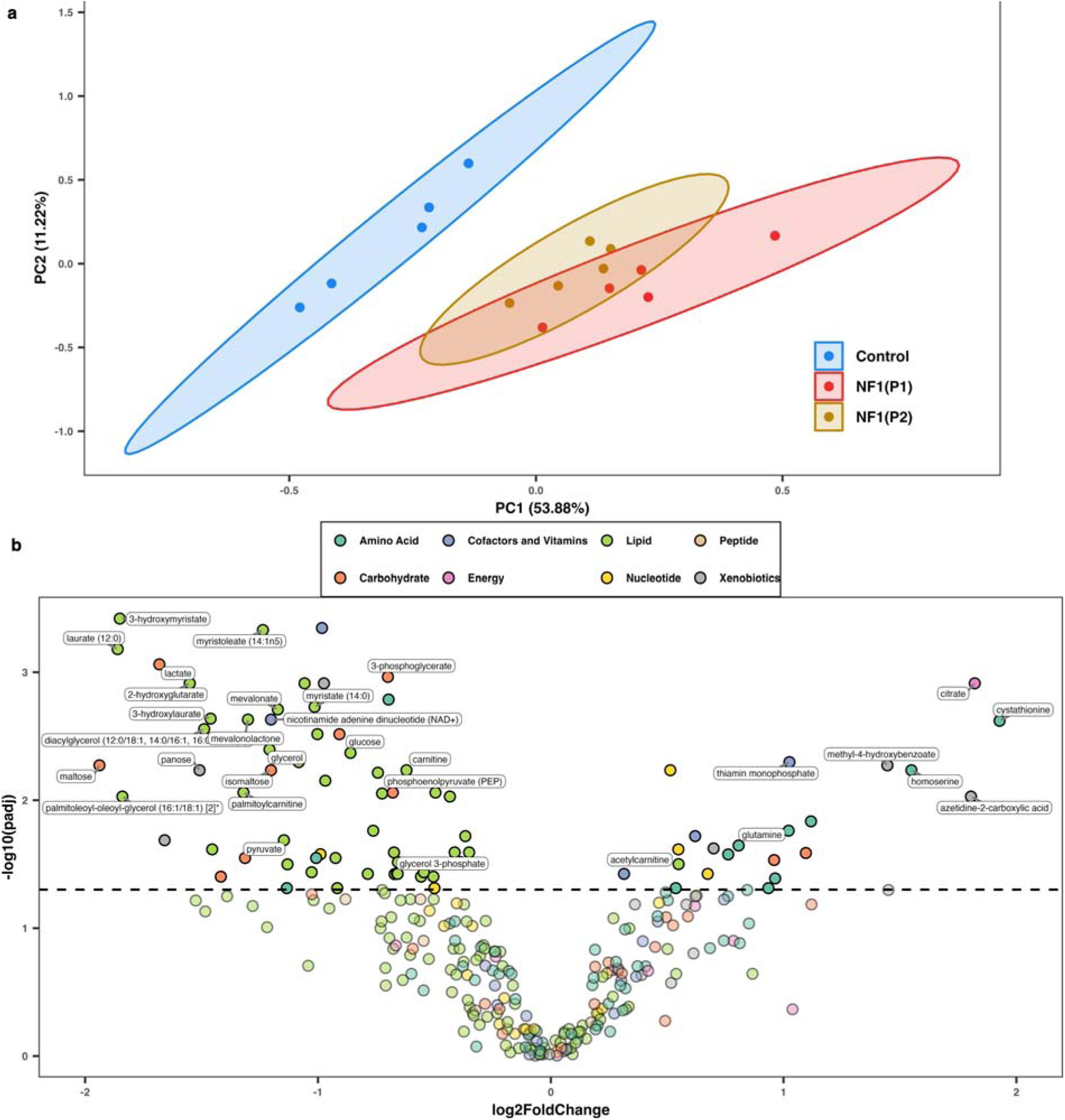
Metabolic profile of *Nf1*-null mutants shows distinct separation from age-matched controls. **(a)** PCA plot of data obtained from an unbiased metabolomic screen, demonstrating distinct separation between the metabolome of two independent *Nf1*-null mutants and age-matched controls. Shaded elliptical regions represent 95% confidence intervals. Each point represents a biological replicate (n= 5). **(b)** Volcano plot illustrating differential metabolite abundance in the *Nf1 Drosophila* model. Each point represents a metabolite, with the x-axis indicating the log2 fold change in metabolite concentration between the average of *Nf1*-null animals (P1 and P2) and age-matched controls, and the y-axis representing the statistical significance (-log10 p-value) of the differential expression. All points above the dashed line (p=.05) denote metabolites that are significantly up- or down-regulated in *Nf1*-null mutants

### Metabolic and behavioral phenotypes are linked in *Nf1*-null *Drosophila*

Though *Nf1* mutant *Drosophila* exhibit a metabolic phenotype indicative of a generalized dearth of energy metabolites, we and others find that these animals exhibit excessive feeding behavior(*17*) (Fig. 2d). In addition, we found nearly every *Nf1*-null animal assessed to have a dramatically enlarged crop (Fig. 2c), the dipteran organ analogous to the human stomach. As feeding is a homeostatic behavior that is tightly regulated in *Drosophila*, flies allowed to feed *ad-libitum* (as all of the animals in this study have been) do not typically develop engorged crops(*18*). However, crop engorgement is observed in animals that have been starved, then allowed to refeed (*19*). For this reason, we hypothesized that *Nf1*-null *Drosophila* may be experiencing a starvation-like state. Indeed, we observed that both *Nf1*-null mutants exhibit increased transcript levels of the dipteran “starvation hormone”, Adipokinetic Hormone (AKH) (Fig. 3e). In addition, we found that the acetylcarnitine(C2) to carnitine(C0) ratio, which increases during fasting (*20*), was significantly increased in *Nf1*-null *Drosophila* (Fig. 2f). Levels of propionylcarnitine (C3) and isobutyrylcarnitine (C4), short-chain acylcarnitines directly derived from branched-chain amino acid catabolism, increase during prolonged fasting due to the breakdown of branched chain amino acids (*21*, *22*). Though levels of C3 and C4 were under the limit of detection for control animals in our metabolomic analysis, C3 was detected in 4/5 *Nf1^P1^*samples and 2/5 *Nf1^P2^* samples (Extended Data Fig. 2e), while C4 was detected in all *Nf1^P1^* samples and 2/5 *Nf1^P2^*samples (Extended Data Fig.2 f). These findings are consistent with the notion that *Nf1*-null *Drosophila* are experiencing a state of starvation despite supranormal food consumption.

**Figure 2.**
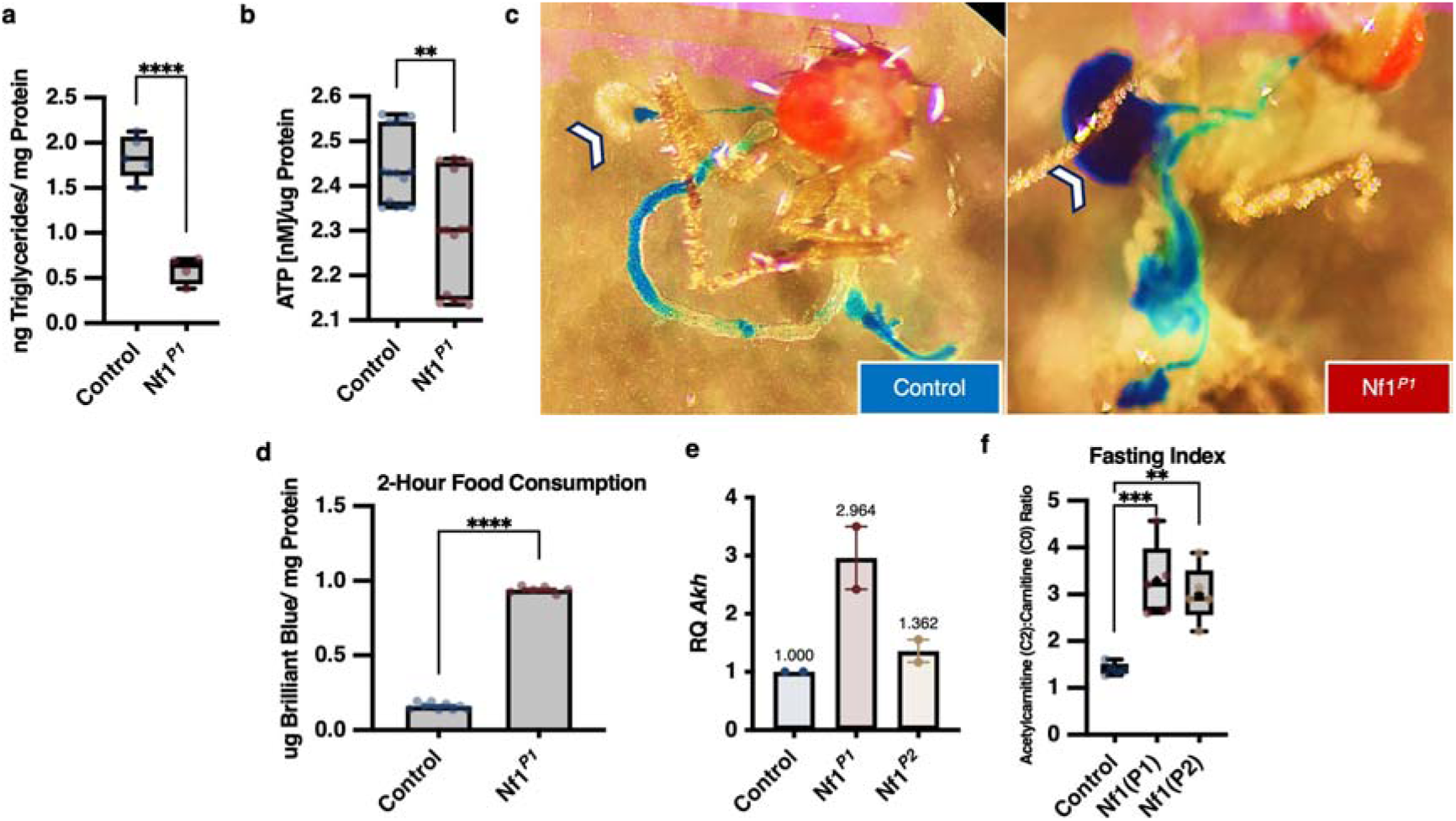
Metabolic profile of *Nf1*-null mutants suggests severe energy deficit. **(a)** Triglyceride levels are substantially reduced in *Nf1*-null mutants (n=5 biological replicates, t(26)=4.764, P<0.0001). **(b)** ATP levels are significantly decreased (n=3 biological replicates, t(28)=3.631, P= 0.0011). **(c)** Loss of *Nf1* leads to exaggerated distension of the *Drosophila* crop (white arrow). **(d)** Loss of *Nf1* results in excessive feeding behavior (n=2 biological replicates, t(13)=66.22, P<0.0001). Data are means±SEM **(e)** *Nf1*-null animals exhibit increased RNA levels of the *Drosophila* “starvation hormone” AKH (n=2 biological replicates). Data are means±SEM. **(f)** Ratio of acetylcarnitine to free carnitine is significantly increased in *Nf1*-null mutants (n=5 biological replicates) One-way ANOVA and Tukey’s multiple comparisons test (F(2, 12) = 15.54, P=0.0005, padj =0.0006, 0.0023). Comparisons are made with unpaired two-tailed t-tests (a,b,d). Box and whisker plots show median, 25th and 75th percentiles, and minimum and maximum values (a,b,f). * P<0.05, ** P<0.01, **** P<0.0001

**Figure 3.**
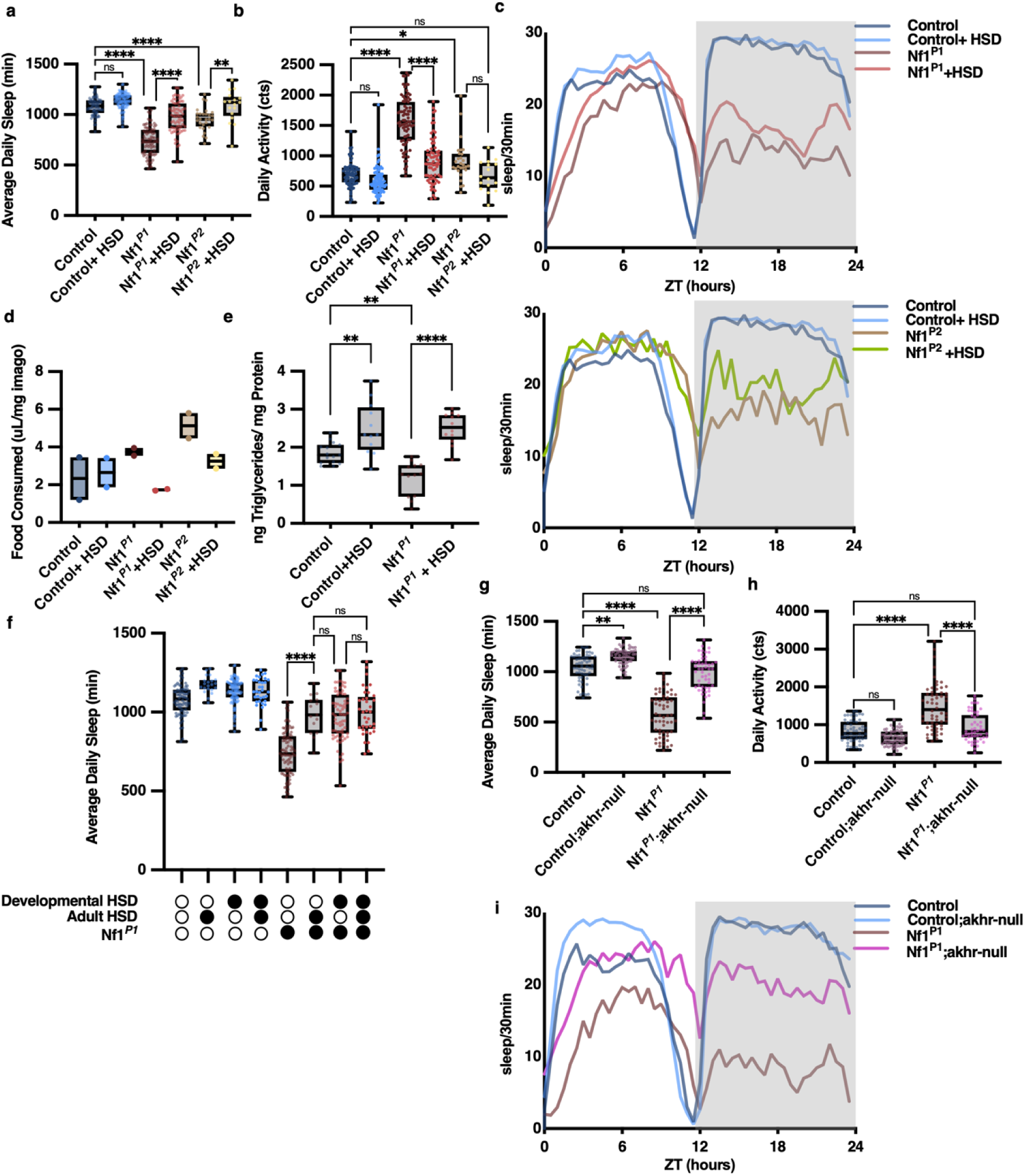
A high sugar diet or decrease in AKH signaling rescues behavioral phenotypes of *Nf1* mutants. **(a)** High sucrose diet (HSD) rescues sleep loss in *Nf1*-null animals (n= 72,94,82,71,28,17[L to R], F(5, 358) = 109.1, P<0.0001). **(b)** *Nf1*-null animals exhibit increased locomotor activity compared to controls, which is attenuated when supplemented with HSD (n= 72,94,82,71,28,17[L to R], F(5, 358) = 103.4, P<0.0001). **(c)** Averaged sleep plots of *Nf1*-null animals and age-matched controls on standard diet and HSD on the second day of the experiment. **(d)** Administration of HSD attenuates excessive feeding behavior in *Nf1*-null animals (n= 2 biological replicates). **(e)** Administration of HSD restores lipid levels (n= 2 biological replicates, F(3, 48) = 22.90,P<0.0001). **(f)** Supplementation of HSD during adulthood is suLicient to rescue of sleep loss in *Nf1* mutants (n = 72, 21, 94, 33, 82, 15, 71, 37[L to R], F(5, 358) = 109.1, P<0.0001). **(g)** Sleep duration is enhanced in *Nf1*^P1^mutants lacking the *akhr* gene (n=57,59,58,51[L to R], F(3, 221) = 152.5, P<0.0001). **(h)** Activity levels are significantly attenuated in *Nf1*^P1^mutants lacking *akhr* as compared to *Nf1* mutants (n=57,59,58,51[L to R], F(3, 221) = 47.75, P<0.0001). **(i)** Averaged sleep plots of *Nf1*^P1^, *akhr-*null mutants with age-matched controls on the second day of the experiment. Box and whisker plots show median, 25th and 75th percentiles, and minimum and maximum values. Comparisons are made with one-way ANOVA and Šídák’s multiple comparisons test. * P<0.05, ** P<0.01, **** P<0.0001.

In *Drosophila*, starvation triggers a neurohormonal cascade that begins with the release of AKH and results in loss of sleep and locomotor hyperactivity (*23*). Interestingly, *Nf1*-null *Drosophila* exhibit reduced sleep, which phenocopies the sleep phenotype exhibited by patients with NF1(*4*, *16*, *24*). In addition, *Nf1*-null *Drosophila* exhibit hyperactivity, which phenocopies the predisposition to ADHD exhibited by patients with NF1(*8*, *25*). We therefore hypothesized that the metabolic phenotype of *Nf1*- null *Drosophila* may contribute to these behavioral phenotypes.

We reasoned that if the sleep phenotype is caused by starvation, then a bolus of easily digested food might mitigate the phenotype. In fact, we found that a high-sucrose diet (HSD) rescued sleep behavior of *Nf1*-null *Drosophila* measured using standard single-beam actigraphy (Fig. 3a, c), as well as multi-beam actigraphy (Extended Data Fig.3d,e), which allows for high resolution determination of sleep: wake behavior. We observed that the HSD rescued sleep behavior even when the animals were only exposed to the diet during the experimental period (Fig. 3a, f, Extended Data Fig. 3g). The high-sucrose diet also suppressed hyperactivity in these animals (Fig. 3b), Extended Data Fig. 3f). HSD did not rescue abnormalities in sleep architecture in *Nf1*-null *Drosophila* (Extended Data Fig. 3 a, b) nor did it rescue circadian rhythmicity (Extended Data Fig. 3c), suggesting that the abnormalities in circadian rhythmicity and sleep architecture exhibited by *Nf1*-null *Drosophila* are regulated by a distinct mechanism. We next tested whether the high-sucrose diet could rescue the metabolic phenotype of *Nf1*-null *Drosophila* and found that the HSD rescued the hypolipidemia observed in *Nf1*-null *Drosophila* (Fig. 3e) as well as the feeding behavior of these animals (Fig. 3d). Rescue by high sucrose food supported our hypothesis that *Nf1* flies chronically experience a state of starvation.

To assess whether the loss of sleep in *Nf1*-null Drosophila is secondary to starvation-induced AKH activity, we attempted to rescue behavioral parameters in *Nf1*-null *Drosophila* by manipulating the AKH pathway. We found that genetic reduction of the AKH pathway by introducing a loss-of-function allele of the AKH receptor (*akhr*) rescued the loss of sleep and hyperactivity in *Nf1*-null *Drosophila* (Fig. 3g, h , Extended Data Fig. 3i,j). These results indicate that loss of *Nf1* causes an energy crisis, leading to the loss of sleep and hyperactivity, the typical behavioral sequelae of starvation in *Drosophila*.

### Non-neuronal Loss of *Nf1* is sufficient to cause Metabolic and Sleep Loss Phenotypes

Much of the work that has been done to understand the etiologies of phenotypes in NF1 has focused on the role of *Nf1* in the nervous system. However, given that energetic phenotypes are just as likely to arise outside the nervous system, we asked if the starvation-induced sleep loss could be recapitulated by knocking down *Nf1* in the periphery. *Nf1* is ubiquitously expressed in *Drosophila*; in mammals, its expression is highest in metabolically active tissues, including muscle, skin, and brain (*26*).

Furthermore, loss of *Nf1* increases energy expenditure in both *Drosophila* and humans (*15*, *17*). The clinical severity of NF1 has not been linked to specific mutations but is negatively correlated with the amount of *Nf1* mRNA (*27*), based upon which we hypothesized that the starvation phenotype observed in *Nf1* mutants is caused by a magnitude-dependent loss of *Nf1* function in highly metabolically active tissues.

To drive *Nf1* knockdown in a tissue-restricted manner, we expressed *Nf1* RNAi with the C179 driver, which drives expression exclusively in mesoderm (*28*) and *mef2*, which is expressed in striated muscle (*29*) (Fig.4a). As noted in other studies, we found that knockdown of *Nf1* using the pan neuronal driver, nsyb, is sufficient to cause metabolic and behavioral phenotypes in mutant *Drosophila* (*16*, *17*, *30*) (Fig. 4b). Though C179 and *mef2* share overlapping expression patterns with *Nf1*, they are not expressed in neurons and share minimal expression overlap with nsyb (Extended Data Fig. 4a). We found that expression of *Nf1* RNAi driven by C179 and mef2 also caused sleep loss and hyperactivity in the flies (Fig.4 c, d). Knockdown of *Nf1* with myoDF31, which is also mesodermally expressed, but in a much more restricted manner chiefly in the digestive system(*31*), does not disrupt the sleep of the animals (Fig. 4e). The finding that broad knockdown of *Nf1* in different (nonoverlapping) tissues produces a similar sleep phenotype supports the previous finding that the NF1 phenotype is negatively correlated with the magnitude of *Nf1* expression (*27*). Finally, we found that either pan-neuronal or pan-muscular knockdown of *Nf1* was sufficient to cause an increase in the transcriptional levels of *Akh* (Fig. 4f). These results suggest that *Nf1* is required for the regulation of energy expenditure in the periphery and the brain, and that the behavioral and metabolic phenotypes in *Nf1* mutants can be caused by magnitude-dependent loss of *Nf1* in either region.

**Figure 4.**
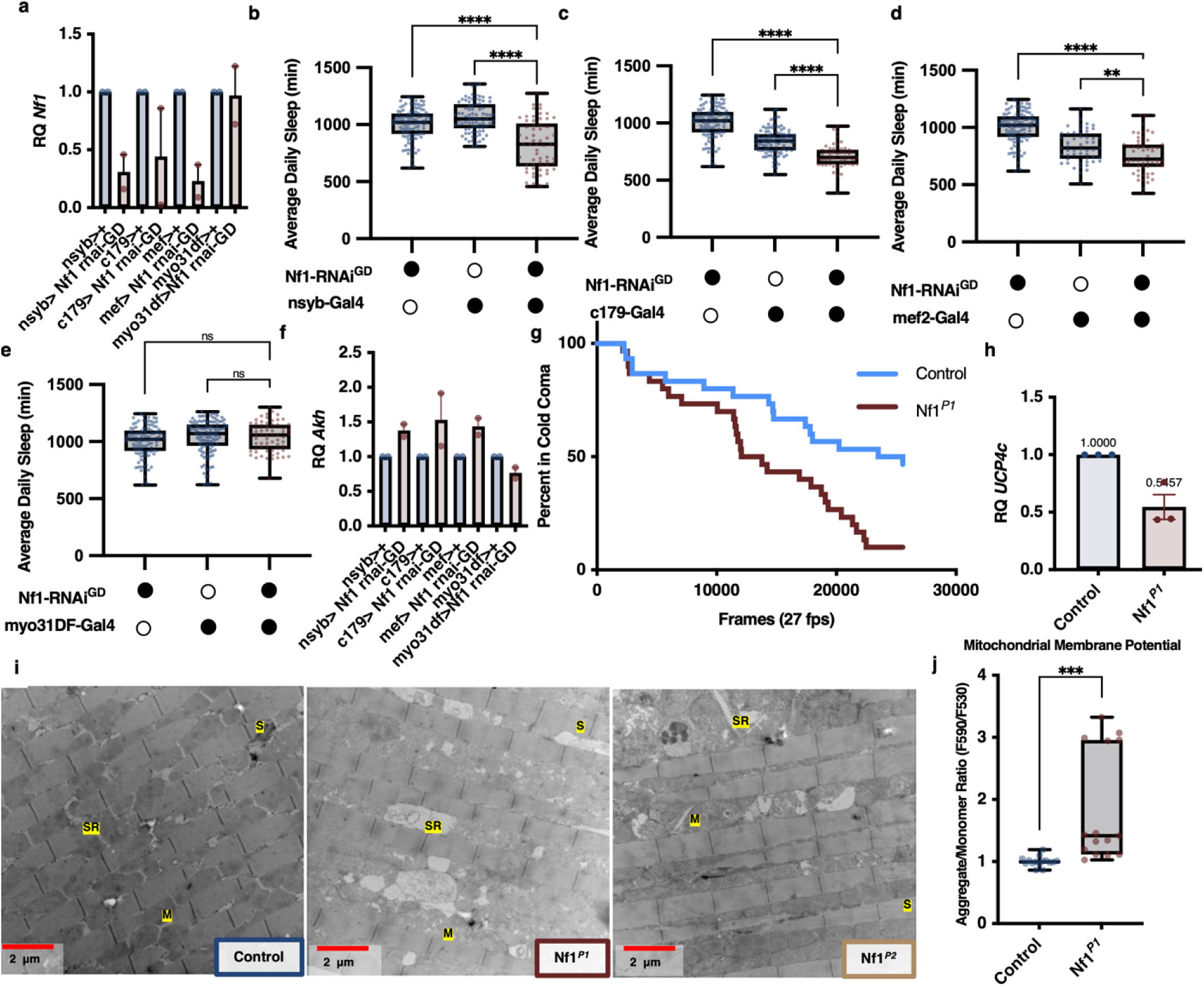
Constellation of metabolic abnormalities in *Nf1* mutants suggests defects in metabolism, especially mitochondrial metabolism, are the source of *Nf1*-null metabolic crisis. **(a)** qPCR results showing that targeted knockdown of *Nf1* in muscle tissue indeed results in a significant reduction of *Nf1* expression (n= 2 biological replicates). **(b)** Neuronal knockdown of *Nf1* results in sleep disturbance, as evidenced by reduced sleep duration (n = 112, 85, 62[L to R]). **(c)** Mesoderm-specific knockdown of *Nf1* leads to sleep disturbance, indicating the involvement of *Nf1* expression in mesodermal tissues in sleep regulation (n= 112,104, 51[L to R]). **(d)**Muscle-specific knockdown of *Nf1* significantly reduces sleep duration, suggesting the involvement of muscle tissue metabolism in sleep homeostasis (n= 112,53,56[L to R]). **(e)** Sleep duration remains unaffected when using a more restricted mesoderm driver for *Nf1* knockdown (n= 112,120,66[L to R]). Comparisons are made with one-way ANOVA and Šídák’s multiple comparisons test (F(8, 700) = 64.25, P<0.0001)(b-e). **(f)** Muscle-specific knockdown of *Nf1* induces increased expression of *Akh* mRNA, indicating a potential mechanism linking *Nf1* deficiency, metabolic regulation, and sleep disturbance (n= 2 biological replicates). (g) *Nf1*-null animals exhibit faster recovery from cold coma. Representative traces of recovery from cold coma after 1hr at 4°C (n= 30). Log-rank (Mantel-Cox) Test χ^2^(1) = 9.200, *p* = 0.0024 and Gehan-Breslow-Wilcoxon Test χ^2^(1) = 6.134, *p* = 0.0133. (h) *UCP4* mRNA expression is reduced in *Nf1*-null animals (n= 3 biological replicates). Data are presented as mean±SEM. **(i)** Representative electron micrographs of *Drosophila* indirect flight muscle at 20,000× magnification. Scale bar indicates 2 microns. Mitochondria (M), sarcomere (S), sarcoplasmic reticulum (SR). **(j)** Mitochondrial membrane potential is elevated in *Nf1*-null mutant mitochondria (n=3 biological replicates, Two-tailed t-test, t(28) = 3.688, P=0.0010). Box and whisker plots show median, 25th and 75th percentiles, and minimum and maximum values (b-e, j). * P<0.05, ** P<0.01, **** P<0.0001.

### Effects of *Nf1* Loss on Energy Expenditure and Mitochondrial Function

Our results so far strongly suggest that loss of NF1 results in an increase in energy expenditure so dramatic that ultimately the whole organism experiences a state of physiological starvation. However, the mechanisms for this increase in energy consumption are not yet clear. In our metabolomic dataset, we noticed that although the overwhelming majority of lipid species were reduced in *Nf1*-null mutants (Extended Data Fig. 2a), a small number of lipid species were elevated. In particular, we found that levels of free fatty acids with a carbon length >14 were normal or sometimes significantly elevated in *Nf1*-null mutants, while fatty acids with a carbon length <14 were significantly reduced (Extended Data Fig. 2a, b). This finding is not only consistent with the idea that *Nf1*-null mutants are undergoing a state of physiological starvation, but also, because efficient catabolism of fatty acids with a carbon length >14 requires functional mitochondrial membrane transport, the accumulation of such fatty acids suggests that mitochondrial function may be altered in *Nf1*-null mutants. Several additional astonishing pieces of data from our studies suggest that the metabolic crisis observed in *Nf1*-null mutants is caused by mitochondrial dysfunction. In particular, we found that *Nf1*-null animals recover more quickly from cold coma (Fig. 4g), which suggests that *Nf1*-null animals are able to generate more heat than controls, consistent with the hypothesis that *Nf1* animals undergo wasteful energy expenditure. Indeed, loss of *Nf1* increases energy expenditure in both Drosophila and humans (*15*, *17*). However, heat generation is unlikely to be UCP (uncoupling protein)-mediated, as we found the mRNA of UCP4C, the UCP most closely linked to increased heat production in *Drosophila* (*32*), to be reduced in *Nf1* animals (Fig. 4h). To investigate the morphology of mitochondria in *Nf1*-null mutants, we performed transmission electron microscopy (TEM) on the indirect flight muscle. In the indirect flight muscle of *Drosophila*, mitochondria are arranged in a regular pattern of rows that are parallel to the muscle fiber, thus differences in mitochondrial histology can be readily observed due to the orderly and stereotyped arrangement of the tissue. In *Nf1*-null mutants, we found many irregularities in the tissue, including the presence of large, swollen mitochondria and abnormally large sarcoplasmic reticulum (Fig. 4i, Extended Data Fig. 5). The swelling of these organelles suggests Ca2+ overload, which is known to cause mitochondrial dysfunction and heat generation (*33*). Indeed, we previously showed that cytoplasmic calcium is elevated in both *Drosophila* and mammalian cells lacking NF1 (*34*). Excess Ca2+ may explain the supranormal heat generation we observed in *Nf1*-null mutants (Fig. 4g).

To investigate mitochondrial function in *Nf1*-null mutants, we measured mitochondrial membrane potential (MMP) in mitochondria isolated from *Nf1*-null mutants and controls. The MMP is a measure of the electrochemical gradient across the inner mitochondrial membrane and is generated by the electron transport chain. We found MMP was dramatically elevated in mitochondria isolated from *Nf1*-null mutants (Fig. 4j). Although this finding could indicate increased mitochondrial activity, in the context of the histological and behavioral data presented above, we believe it to be more likely that elevated MMP is due to mitochondrial dysfunction secondary to calcium overload, as the capacity for mitochondrial Ca2+ uptake is dictated by the MMP. Hyperpolarized MMP is necessary for the uptake of excess Ca2+ into the mitochondrial matrix (*35*). Thus, the elevated MMP observed in *Nf1*-null mutants is likely a compensatory response to Ca2+ overload. Taken together, these results provide evidence for the role of mitochondrial dysfunction as a potential underlying mechanism for the excessive energy utilization and subsequent metabolic crisis observed in *Nf1*-null mutants.

### SARM1 hyperactivity causes NAD+ depletion and sleep loss in *Nf1*-null mutants

The metabolic phenotypes observed in *Nf1*-null mutants suggest that *Nf1* is required for the regulation of energy expenditure. However, the mechanism by which *Nf1* regulates energy expenditure, or why its loss increases calcium, is unknown. To investigate this, we looked to the data from our metabolomic analysis of *Nf1*-null mutants (Fig. 1) and noticed that *Nf1*-null mutants have a dramatic depletion of Nicotinamide Adenine Dinucleotide (NAD+) and related metabolites (Extended Data Fig. 1 h-k). Our independent determination of NAD+ levels in *Nf1*-null animals concurred with this finding (Fig. 5a). NAD+ is a coenzyme required for the function of many metabolic enzymes and mitochondrial function, suggesting that its depletion is related to the mitochondrial dysfunction observed in *Nf1*-null mutants. Supplementation with nicotinamide (NAM), an NAD+ precursor, can rescue NAD+ levels depleted by mitochondrial dysfunction in *Drosophila* (*36*). NAM rescues NAD+ levels by two mechanisms: by providing a precursor for NAD+ synthesis, and by product inhibition of NAD+ consuming enzymes, such as PARPs, sirtuins, CD38, and SARM1 (*37*). Consistent with a role for NAD+ depletion in the metabolic crisis that causes sleep behavior dysregulation in *Nf1*-null *Drosophila*, NAM supplementation rescued sleep loss and attenuated hyperactivity in *Nf1* animals while under both single beam and high resolution multibeam surveillance (Fig.5 c, d, Extended Data Fig.6 a, c). Partial rescue may reflect an inability to completely repair damage already done.

**Figure 5.**
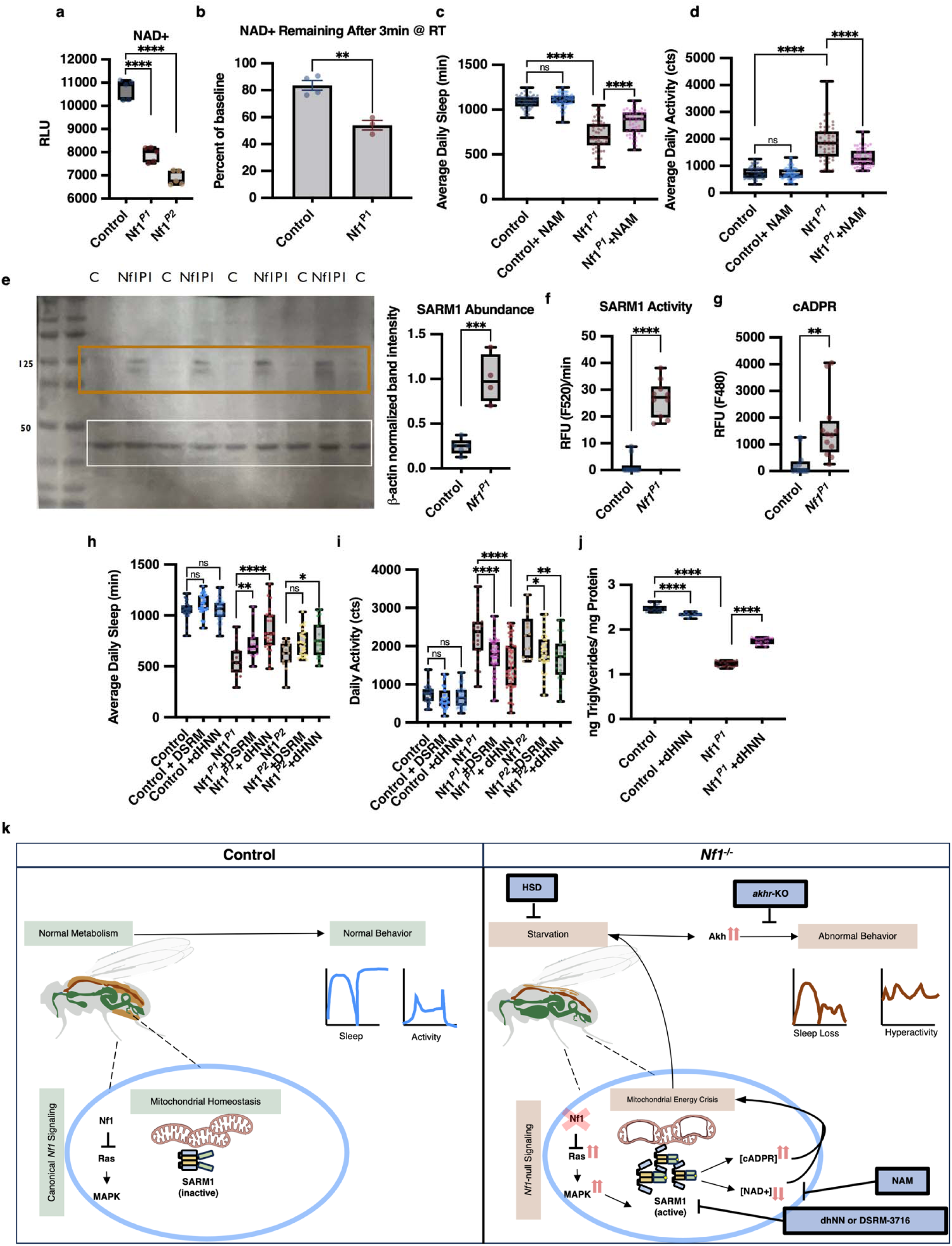
SARM1 hyperactivation is responsible for metabolic crisis and sleep loss in *Nf1*-null mutants. **(a)** *Nf1*-null mutants display significantly reduced levels of NAD+ (n= 5 biological replicates, F(2, 12) = 158.8, P<0.0001). **(b)** Addition of NAD+ to *Nf1*-null lysate results in rapid depletion. Quantification of NAD+ depletion in *Nf1*-null lysate 3 min after [NAD+] spiked to 100uM NAD+ (n = 3-4 technical replicates, Two-tailed t-test, t(5) = 5.711, P=0.0023). Data are means±SEM. **(c)** Supplementation with 10 mM Nicotinamide (NAM), an NAD+ precursor, rescues sleep loss in *Nf1*-null animals (n= 53,62,50,52[L to R], F(3, 213) = 148.7, P<0.0001). **(d)** Supplementation with 10 mM NAM attenuates hyperactivity in *Nf1*-null animals (n= 53,62,50,52[L to R], F(3, = 105.5, P<0.0001).**(e)** Representative blot and densitometric analysis illustrate the upregulation of SARM1protein expression in *Nf1*-null animals. Orange box = SARM1 (MW = 130 kDa), white box = B-actin (MW = 42 kDa) Quantification of SARM1 protein expression in *Nf1*-null animals normalized to loading control (n= 4-5 lanes, two-tailed t-test, t(7) = 5.879, P=0.0006). **(f)** Measurement of SARM1 NADase activity in *Nf1*-null tissue shows SARM1 specific enzymatic conversion of PC6 to fluorescent PAD6 is significantly increased compared to controls, indicating SARM1 hyperactivation in *Nf1*-null tissue (n= 2 biological replicates, two-tailed t-test, t(18) =10.59, P<0.0001). **(g)** Quantification of cADPR levels in *Nf1*-null lysate shows a significant elevation compared to controls (n= 2 biological replicates, two-tailed t-test, t(22) = 3.504, P=0.0020). **(h)** Pharmacological inhibition of SARM1 with 10uM dHNN or 50uM DSRM corrects sleep behavior in *Nf1*-null mutants (n= 31,12,30,8,31,10[L to R], F(3, 213) = 148.7). **(i)** Pharmacological inhibition of SARM1 corrects locomotor hyperactivity in *Nf1*-null mutants (n= 31,12,30,8,31,10[L to R], F(3, 213) = 148.7, P<0.0001). **(j)** Pharmacological inhibition of SARM1 with 10uM dHNN for seven days restores triglyceride levels in *Nf1*-null mutants (n=4-5 biological replicates, F(3, 71) = 1523, P<0.0001. Box and whisker plots show median, 25th and 75th percentiles, and minimum and maximum values. Comparisons are made with one-way ANOVA and Šídák’s multiple comparisons test. For panels b, d, e, and f, comparisons are made with unpaired two-tailed t-tests. * P<0.05, ** P<0.01, **** P<0.0001 **(k)** Overview schematic of proposed model. Loss of Nf1 leads to SARM1 activation, which in turn leads to energetic crisis and mitochondrial dysfunction through depletion of NAD+. AKH signaling is increased as a compensatory response to energetic crisis. Increased AKH signaling leads to hyperactivity and sleep loss. The different manipulations highlighted in blue boxes can block effects of the mutant at different points and rescue sleep levels.

We next sought to investigate how NAD+ becomes so depleted in *Nf1*-null mutants, and so we measured NAD+ levels in *Nf1*- null and control lysates after they were spiked with a standard concentration of NAD+. The NAD+ standard is much more rapidly depleted upon addition to *Nf1*-null lysate compared to control lysate (Fig. 5b). This finding suggests the presence of an aberrantly active NAD+ hydrolase in *Nf1*-null tissue. We considered a role for Sterile Alpha and TIR Motif Containing 1 (SARM1) because it is a recently characterized NAD+ hydrolase that is expressed in mitochondria and activated by the Mitogen-Activated Protein Kinase (MAPK) signaling pathway (*38*), the pathway that is canonically activated by loss of NF1 through upregulation of Ras activity in mammals and in *Drosophila* (*39*). Interestingly, SARM1 hydrolyzes NAD+ to produce cyclic ADP-ribose (cADPR) (*40*), which regulates intracellular calcium levels. Specifically, cADPR is a potent activator of the ryanodine receptor, an intracellular calcium channel expressed in the endoplasmic reticulum of various excitable animal tissue like muscles and neurons (*41*). Thus, calcium release from the endo-/sarco-plasmic reticulum is a downstream effect of SARM1 activation. Calcium is a well-known metabolic accelerant, inducer of thermogenesis, and mitochondrial toxin (at high levels) (*33*), and, as noted above, its concentration increased in *Nf1* mutants. To determine if SARM1 is the NAD+ hydrolase responsible for the mitochondrial dysfunction and metabolic crisis observed in *Nf1*-null mutants, we assayed expression of SARM1 protein and found that it is dramatically increased in *Nf1* mutants (Fig. 5e), as is SARM1-specific NADase activity (*42*) (Fig. 5f, Extended Data Fig. 6b). Finally, because SARM1 preferentially produces cADPR during NAD+ hydrolysis, cADPR is a valuable dosage-sensitive biomarker for SARM1 activity (*40*). cADPR levels are significantly increased in *Nf1*-null lysate (Fig. 5g).

Finally, we asked if pharmacological inhibition of SARM1 could correct sleep behavior in *Nf1*-null mutants. We found that pharmacological inhibition of SARM1 with either the irreversible inhibitor dehydronitrosonisoldipine (dHNN) or the reversible inhibitor 5-Iodoisoquinoline (DSRM-3716) increased sleep (Fig. 5h) and triglyceride levels in *Nf1*-null mutants (Fig. 5i), strongly suggesting that SARM1 hyperactivation contributes to metabolic crisis and sleep loss we have observed in *Nf1*-null mutants. Taken together, these results provide evidence for the critical role of mitochondrial dysfunction and SARM1 hyperactivation in the metabolic crisis and sleep loss observed in *Nf1*-null mutants.

### Conclusions and Future Directions

Our findings reveal that a constellation of metabolic defects, including severe energy deficit and perturbations in lipid and carbohydrate metabolism, culminate in a starvation-like state that accounts for much of the sleep loss observed in *Nf1* mutants. Additionally, we discovered that muscle-specific loss of *Nf1* was sufficient to cause these sleep and metabolic phenotypes, indicating that effects of NF1 on cellular energy expenditure in the periphery can impact overall organismal physiology and behavior. We traced the basis of the metabolic crisis, and thereby behavioral phenotypes, observed in *Nf1* mutants to defects in mitochondrial function, and specifically in the activity of the enzyme SARM1. Our findings provide new avenues for investigating the mechanisms underlying the pathophysiology of NF1 and exploring potential therapeutic targets to ameliorate the effects of this debilitating disease.

## Materials and Methods

### Fly genetics and husbandry

Fly strains that contain the *Nf1P1*, *Nf1P2*, and *K33* alleles are described in (*43*). C179-Gal4, Mef2-Gal4, and Myo31DF-Gal4 were obtained from the Bloomington Drosophila Stock Center (BDSC). UAS-NF1 RNAi GD was obtained from the Vienna Drosophila Resource Center (VDRC). All fly strains were typically cultured on a standard cornmeal-molasses medium and maintained in 12 h light: 12 h dark (LD) cycles at 25 °C, except for diet rescued animals, which had been cultured on their indicated diets. Recipes for high sucrose and 10mM NAM diets were adapted from (*44*). High sucrose and 10mM NAM diets are identical to the compared standard diet, save for the addition of the supplement. For the high sucrose and NAM supplementation experiments, flies were raised entirely on the indicated diet, including throughout development, except where explicitly noted. For the SARM1 inhibitor drug treatment experiments, flies were raised on standard cornmeal-molasses food and then transferred to circadian tubes with food containing t 10 uM dHNN or 50uM DSRM only for the duration of the experiment. Only male flies were used in all experiments.

### Metabolomics

5- to 7-day-old adult male flies were collected on dry ice. Flies of each genotype were pooled into five independent groups comprised of 50 flies each. These samples were then biochemically profiled at Metabolon, Inc., as described in (*45*). Raw data were extracted, peak-identified, and QC processed using Metabolon’s proprietary hardware and software. Code and raw data used to generate visualizations in Fig. 1 and Extended Data Fig. 1 can be found at github.com/folasofela/nf1metabolism (repository will be made public on submission).

### Feeding Assays

Food consumption was measured in one of two ways: (1) The CaFE assay (a non-invasive method of measuring food consumption in *Drosophila*) (*46*) was performed on groups of 15 flies per replicate. Ten flies were placed in vials with a moist Kimwipe as a water source and 5% aqueous sucrose available in a calibrated capillary tube. The capillary tube was marked at the beginning and end of the assay and the amount of food consumed was determined by measuring the distance between the marks. 2) Because *Nf1* mutant flies have some difficulty with climbing, we also measured food consumption by assaying the amount of food consumed by a group of flies over time. Briefly, flies were placed in a vial containing a small amount of 5% sucrose, 2% agar food dyed with Brilliant Blue FCF, a non-toxic food dye. After 2 hours, the flies were homogenized in PBS and the amount of food consumed was determined spectrophotometrically by measuring the absorbance of the homogenate at 630nm.

### Sleep and Activity Assays

The *Drosophila* Activity Monitor (DAM) system (TriKinetics, Inc., Waltham, MA) was used to measure locomotion in 5–7-day old flies as in (*47*). Sleep is scored as a period of inactivity lasting at least 5 minutes. Flies were raised throughout development on standard, high sucrose, or 10mM NAM food, and then assayed for sleep in tubes containing 5%sucrose, 2% agar for the duration of the experiment, except where explicitly noted. SARM1 inhibitor drug treatments were given to flies only during the assay by 10 uM dHNN or 50uM DSRM dHNN directly into the agar used to prepare DAM tubes. Flies that died at any time during the assay were excluded from analysis. Raw actigraphy data and the code used to generate sleep and activity plots can be found at github.com/folasofela/nf1metabolism (repository will be made public on submission).

### Biochemical Measurements

5- to 7-day-old adult male flies were collected on dry ice in groups of 10 flies between ZT 6 and 8. The samples were then homogenized in an appropriate preparatory buffer. For absorbance-based measurements of Triglycerides flies were homogenized in PBS + .05% Tween20 as described in (*48*) and the concentration of triglycerides was determined using the Infinity€ Triglycerides Liquid Stable Reagent (# TR22421) according to the manufacturer’s instructions. For luminescence-based measurements of NAD+, cADPR and ATP, flies were homogenized in 0.2N NaOH + 1% DTAB and processed according to the manufacturer’s instructions: NAD+ (Promega # G9071), cADPR (AAT Bioquest # 20305), ATP (Invitrogen # A22066).Where indicated in Figs. 2, 3, and 5 the protein concentration of samples was measured with the Pierce® BCA Protein Assay Kit (Thermo Scientific # 23225) and used to normalize metabolite concentrations.

### Cold Coma Recovery Assay

Flies were placed individually into opaque white 96 well plates, and the plate was placed in the cold room on ice for 60m. The plate was then returned to room temperature and then tracked for cold coma recovery using MARGO, a MATLAB-based, real-time animal tracking suite for custom behavioral experiments (*49*). Custom code to analyze MARGO output for arousal from cold coma and raw data for this assay can be found at github.com/folasofela/nf1metabolism (repository will be made public on submission).

### Mitochondrial Isolation and Membrane Potential Measurements

Mitochondria from 5–7-day old male flies were isolated from 50 whole fly bodies collected on dry ice between ZT 6 and 8 using the Mitochondria Isolation Kit for Tissue (Thermo Scientific # 89874) according to the manufacturer’s instructions. Mitochondrial membrane potential was measured using the JC-1 - Mitochondrial Membrane Potential Assay Kit (Abcam # ab113850) according to the manufacturer’s instructions. Briefly, 10μg of mitochondria were incubated with 1X JC-1 dye for 15m at 37°C. The ratio of red (excitation/emission: 535/595nm) to green (excitation/emission: 485/535nm) fluorescence was used as a measure of mitochondrial membrane potential.

### Reverse transcription-quantitative PCR (RT-qPCR)

For each biological replicate, RNA was extracted from 10 whole fly bodies collected on dry ice between ZT 6 and 8 using SV RNA Total Isolation kit (Promega #Z3101) according to manufacturer’s instructions. One-step RT-qPCR was done using *Power* SYBR Green PCR master mix (Applied Biosystems #4368577). For each gene, 2 or 3 biological replicates were assayed, comprising three to five technical replicates. PCR conditions were: 48°C for 30m, 95°C for 10m, followed by 40 cycles of 95°C for 15s and 60°C for 1m. Melting curve analysis was carried out for all PCR reactions to verify homogeneity of the PCR products and the absence of primer dimers. Samples with multiple significant peaks in the melting curve were excluded from analysis. CT values were normalized via the 2-ΔΔCTmethod (*50*). Equalized amplicons of Actin5C were used as a reference gene to normalize expression via the following primer sets:

*Akh* F -AGACCTCCAACGAAATGCTG R- GTGCTTGCAGTCCAGAAAGAG *Nf1* F- TATCGCGCTACCGCTTCTC R-TTCCAGAGTGGTCAGTATGATGA *UCP4C* F- ACAAACGTCGCTGATCCACTA R-GGAAGACACACGACTCGGC *ACT5C* F-TTGTCTGGGCAAGAGGATCAG R- ACCACTCGCACTTGCACTTTC *Akhr* F- CACACCTCGCTGTCCAATC R -CATCACCTGGCCTCTTCCA

### Western Blot

Protein from 5-7 day old flies was extracted from 10 whole fly bodies collected on dry ice in groups of 10 flies between ZT 6 and 8. 10 whole fly bodies were homogenized in a modified RIPA buffer (50mM Tris-HCl pH 7.4, 150mM NaCl, 1mM EDTA, 0.1% SDS, 1% Triton X-100, 1mM NaF, 1mM Na3VO4, 1mM PMSF, 1X protease inhibitor cocktail (Roche # 11836170001)) and boiled for 5 minutes. Samples were run on a 4-20% gradient SDS-PAGE gel and transferred to a 0.2μm nitrocellulose membrane. Membranes were blocked in Odyssey Blocking Buffer (LI-COR # 927-40000) for 1 hour at room temperature.

Membranes were washed in TBS-T and incubated with primary antibodies overnight at 4°C. Membranes were washed in TBS- T and incubated with secondary antibodies for 1 hour at room temperature. Primary antibodies used were: anti-SARM1 (D2M5I) Rabbit mAb (Cell Signaling Technologies #13022) at 1:1000 and Anti-beta Actin antibody Loading Control (#Abcam ab8224) at 1:20,000. Anti- Rabbit and Anti- Mouse HRP conjugated secondary antibodies were used at 1:10,000. For band visualization, membranes were incubated in DAB substrate (Thermo Scientific #34002) for 5 minutes and imaged. Band intensities were quantified using ImageJ.

### Muscle Electron Microscopy

Indirect flight muscle from Control (K33), Nf1-P1, and Nf1-P2 flies were dissected in PBS and fixed in 2.5% glutaraldehyde in 0.1M sodium cacodylate buffer and subsequently fixed and processed as in (*45*). Ultrathin sections for electron microscopic examination were prepared. Thin sections were stained with lead citrate and examined with a JEOL 1010 electron microscope fitted with a Hamamatsu digital camera and AMT Advantage image capture software.

### SARM1 Activity Assay

As reported in (*51*) , 4-[(E)-2-(4-Ethoxyphenyl)vinyl]pyridine (PC6) is a compound that is enzymatically processed into the highly fluorescent compound PAD6 exclusively in the presence of activated SARM1 and NAD+. *Drosophila* homogenate from 10 male animals was incubated with 100uM PC6 (ChemDiv # SR00-0002) and 100uM NAD+ (Sigma # N0632) and the fluorescence was measured over 2 hours.

## Acknowledgements

Special thanks to Dr. Eliana D. Weisz and Dr. Lei Bai for their preparation and submission of the metabolomic samples used to generate data for in Fig. 1 and Extended Data Fig. 1 and to Dr. Eliana D. Weisz for the preparation and submission of samples for mitochondrial electron micrographs pictured in Fig. 4 and Extended Data Fig.4.

## Author Contributions

F.A.S, A.S, and T.A.J conceived the project. F.A.S and M.L.V conducted experiments. M.L.V conducted data validation experiments. F.A.S developed software and performed data analyses. F.A.S designed the experiments and wrote the manuscript. All authors contributed to manuscript editing. T.A.J and A.S provided study materials, reagents, materials, and funding. A.S supervised all aspects of the work.

## Data Availability

Raw data is available at github.com/folasofela/nf1metabolism . Publicly available scRNAseq data from FlyCellAtlas (https://www.science.org/doi/10.1126/science.abk2432) was used in Extended Data Fig 4. Any other relevant materials or reagents are freely available or may be obtained the corresponding authors upon reasonable request.

## Code Availability

Code used to analyze data is available at github.com/folasofela/nf1metabolism .

---

**Extended Data Figure 1.**
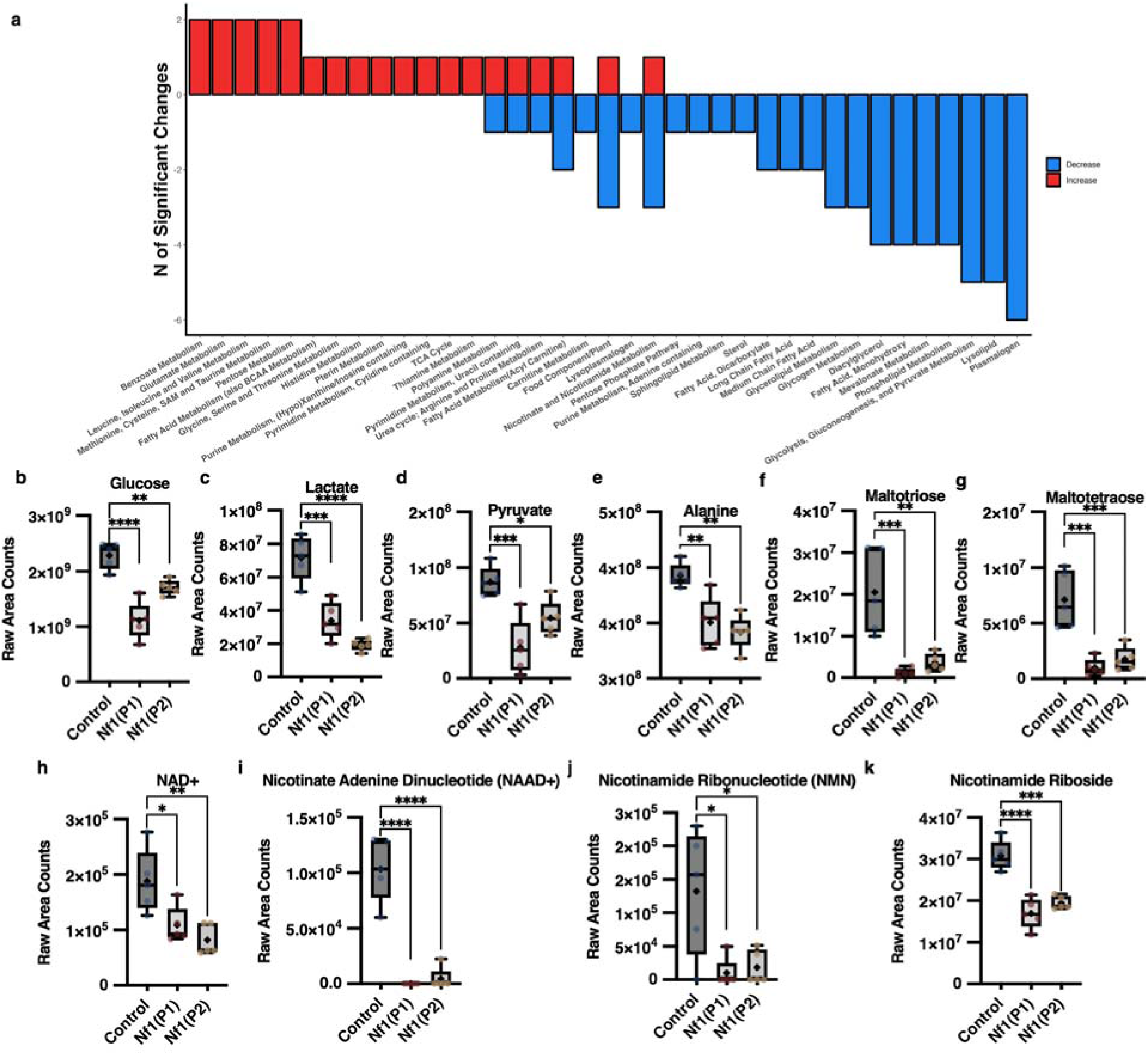
Metabolic profile of *Nf1*-null mutants suggests severe energy deficit. **(a)** Bar plot of number of metabolites in pathways significantly enriched or depleted in both *Nf1*-null mutants (P1 and P2) as compared to age-matched controls. Metabolites were identified using the Kyoto Encyclopedia of Genes and Genomes (KEGG) database. The x-axis represents the number of metabolites in each pathway, and the y-axis represents the pathway name. P value cutoff of 0.05 was used to determine significance. **(b -e)** Box and whisker plots of significantly depleted glycolytic and gluconeogenic metabolites. **(f,g)** Box and whisker plots of significantly depleted TCA cycle metabolites. **(i-k)** Box and whisker plots of significantly depleted NAD+ metabolites. Box and whisker plots show median, 25th and 75th percentiles, and minimum and maximum values. Comparisons are made with one-way ANOVA and Šídák’s multiple comparisons test (b-k). * P<0.05, ** P<0.01, **** P<0.0001.

**Extended Data Figure 2.**
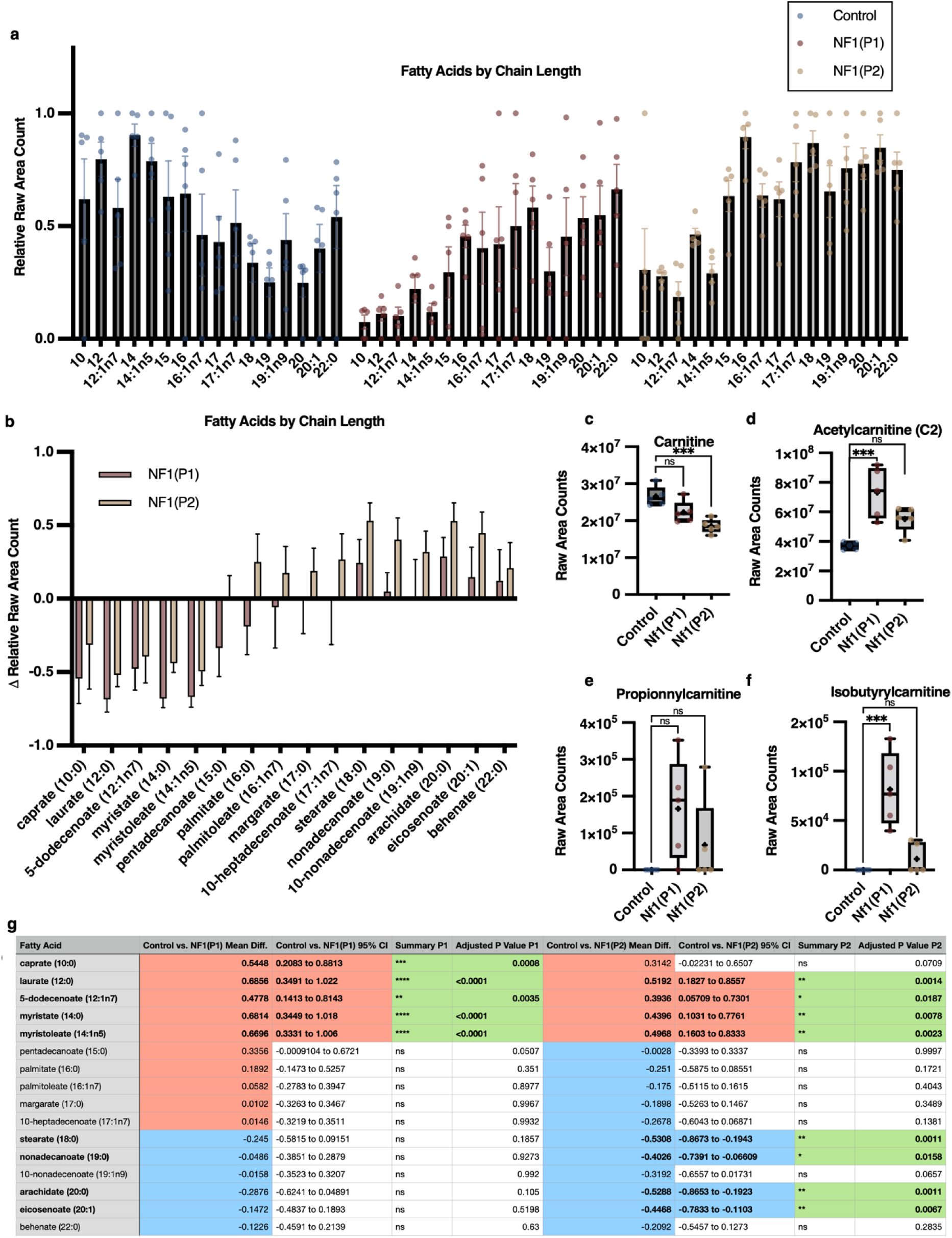
Metabolic profile of *Nf1*-null mutants suggests defects in lipid metabolism. **(a)** Barplot of fatty acids by carbon chain length in *Nf1*-null mutants (P1 and P2) as compared to age-matched controls. Relative raw peak areas were min-max normalized for each metabolite to aid in visualization. **(b)** Barplot of differences in relative fatty acid abundance by carbon chain length in *Nf1*-null mutants (P1 and P2) as compared to age-matched controls. **(c-f)** Box and whisker plots of dysregulated acylcarnitines. Box and whisker plots show median, 25th and 75th percentiles, and minimum and maximum values. comparisons are made with one-way ANOVA and Šídák’s multiple comparisons test (b-f). * P<0.05, ** P<0.01, **** P<0.0001.

**Extended Data Figure 3.**
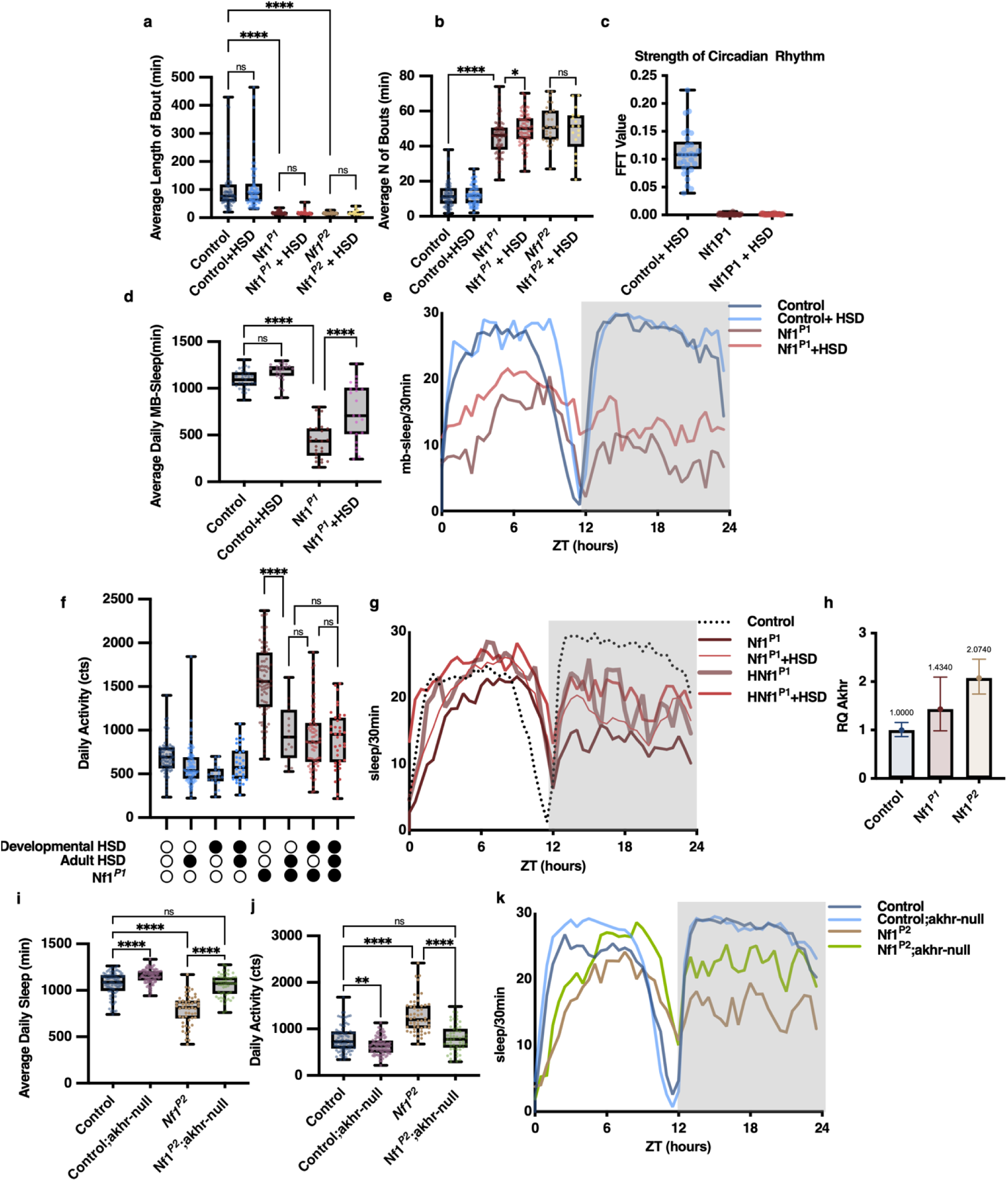
A high sugar diet or decrease in AKH signaling rescues behavioral phenotypes of *Nf1* mutants. **(a-b)** HSD does not significantly improve bout length and bout number, measures of sleep consolidation, in *Nf1*-n€ animals (n = 73, 94, 82,71, 28, 17 [L to R], F (5, 359) = (a) 44.47 (b) 308.7 P<0.0001). **(c)** HSD does not significantly improve strength of circadian rhythms as measured by Fourier Fast Transform (FFT) periodogram analysis in *Nf1*-null animals (n= 29,29,30 [L to R]). **(d)** HSD improves sleep duration in *Nf1*-null animals as measured with multibeam infrared actigraphy (n = 31,24,25,23 [L to R], F(3, 99) = 82.28 P<0.0001).(e) Averaged sleep traces of *Nf1*-null animals on HSD using multibeam infrared actigraphy. **(f)** Supplementation of HSD during adulthood is sufficient to attenuate hyperactivity in *Nf1* mutants (n = 72, 21, 94, 33, 82, 15, 71, 37[L to R], F (7, 418) = 88.52, P<0.0001). **(g)** Averaged sleep traces of *Nf1*-null animals on HSD during adulthood using single beam infrared actigraphy. “H” **=** animals given 25% sucrose in circadian tubes, “+HSD” **=** animals raised on HSD during development **(h)** RNA levels of *Akhr* are significantly increased in *Nf1*-null animals as compared to age-matched controls (n=1 biological replicate). **(i)** Sleep duration is enhanced in *Nf1^P2^* mutants lacking the *akhr* gene (n=82,84,54,54[L to R], F(3, 270) = 73.52, P<0.0001). **(j)** Activity levels are significantly attenuated in *Nf1^P2^*mutants lacking *akhr* as compared to *Nf1* mutants (n=82,84,54,54 [L to R], F(3, 270) = 37.18, P<0.0001). **(k)** Averaged sleep plots of *Nf1^P2^*, *akhr-*null mutants with age-matched controls on the second day of the experiment. Box and whisker plots show median, 25th and 75th percentiles, and minimum and maximum values. Comparisons are made with one-way ANOVA and Šídák’s multiple comparisons test. * P<0.05, ** P<0.01, **** P<0.0001

**Extended Data Figure 4.**
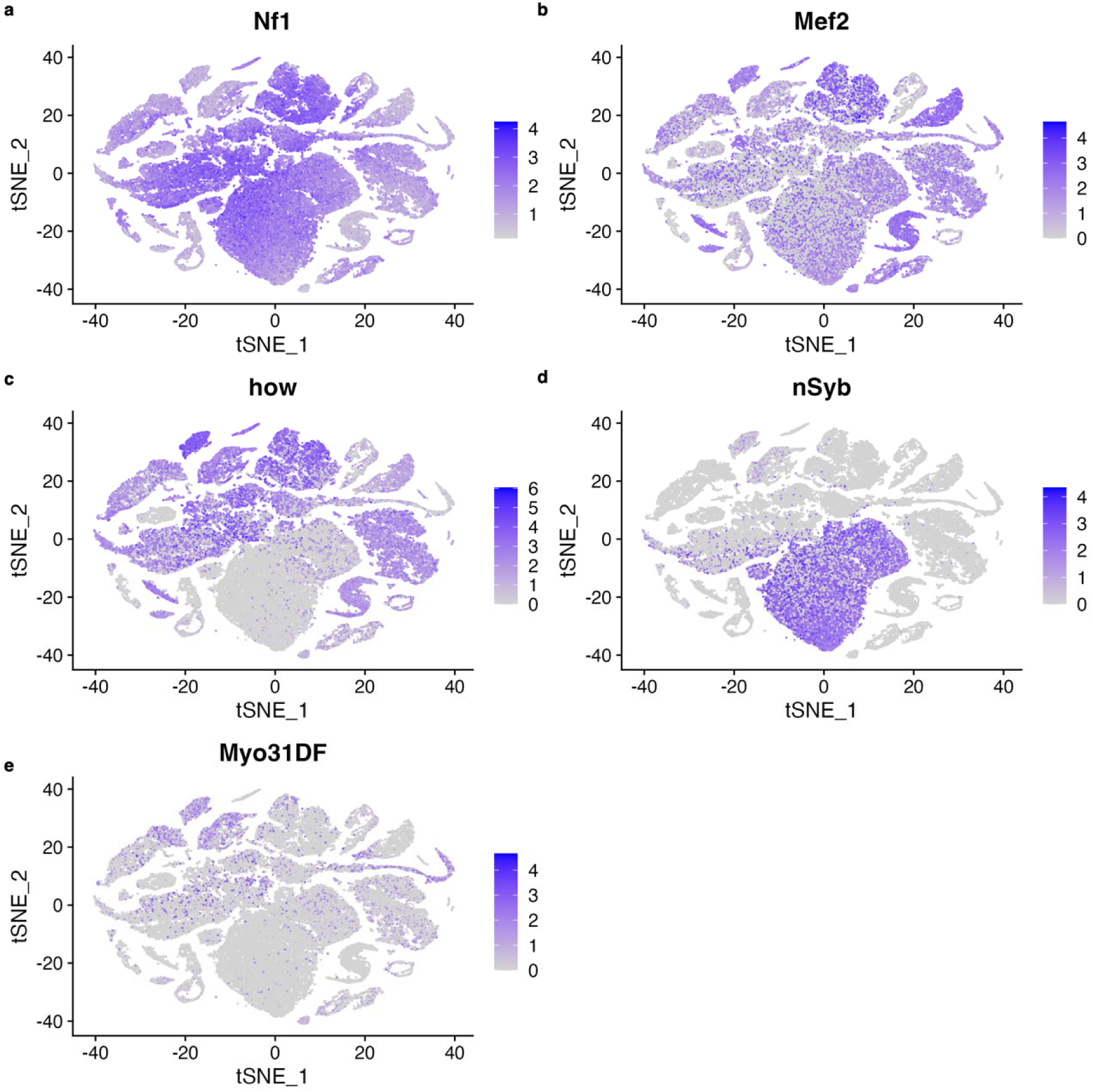
Gene Expression Patterns in Nf1-Positive Cells from Adult Fruit Fly. Unsupervised clustering (Louvain algorithm, modularity 0.9958) of 59,100 Nf1-positive cells from a single-nucleus transcriptomic dataset of 506,660 adult fruit fly cells (Li et al. 2022) identified seven major clusters based on the differential expression of 16366 genes, illustrated via t-SNE. Differential expression patterns of key genes (*Nf1*, *Mef2*, *how*, *nSyb*, and *Myo31DF*) are mapped. *nSyb* expression is prominent in clusters representing neuronal cells, The expression of *how*, which follows the pattern of the enhancer trap line *c179*-gal4, is known to be mesoderm-specific. Plots underscore the lack of overlap between c179-gal4/how and *nSyb* expression. *Mef2* is predominantly in non-neuronal tissues with some scattered neuronal presence. *Myo31DF* expression localizes to mesodermal clusters, but its RNA is sparsely expressed in *Nf1*+ cells compared to *Mef2* and *how,* providing a molecular basis for the absence of a sleep phenotype in Nf1 knockdown experiments using *Myo31DF*-gal4.

**Extended Data Figure 5.**
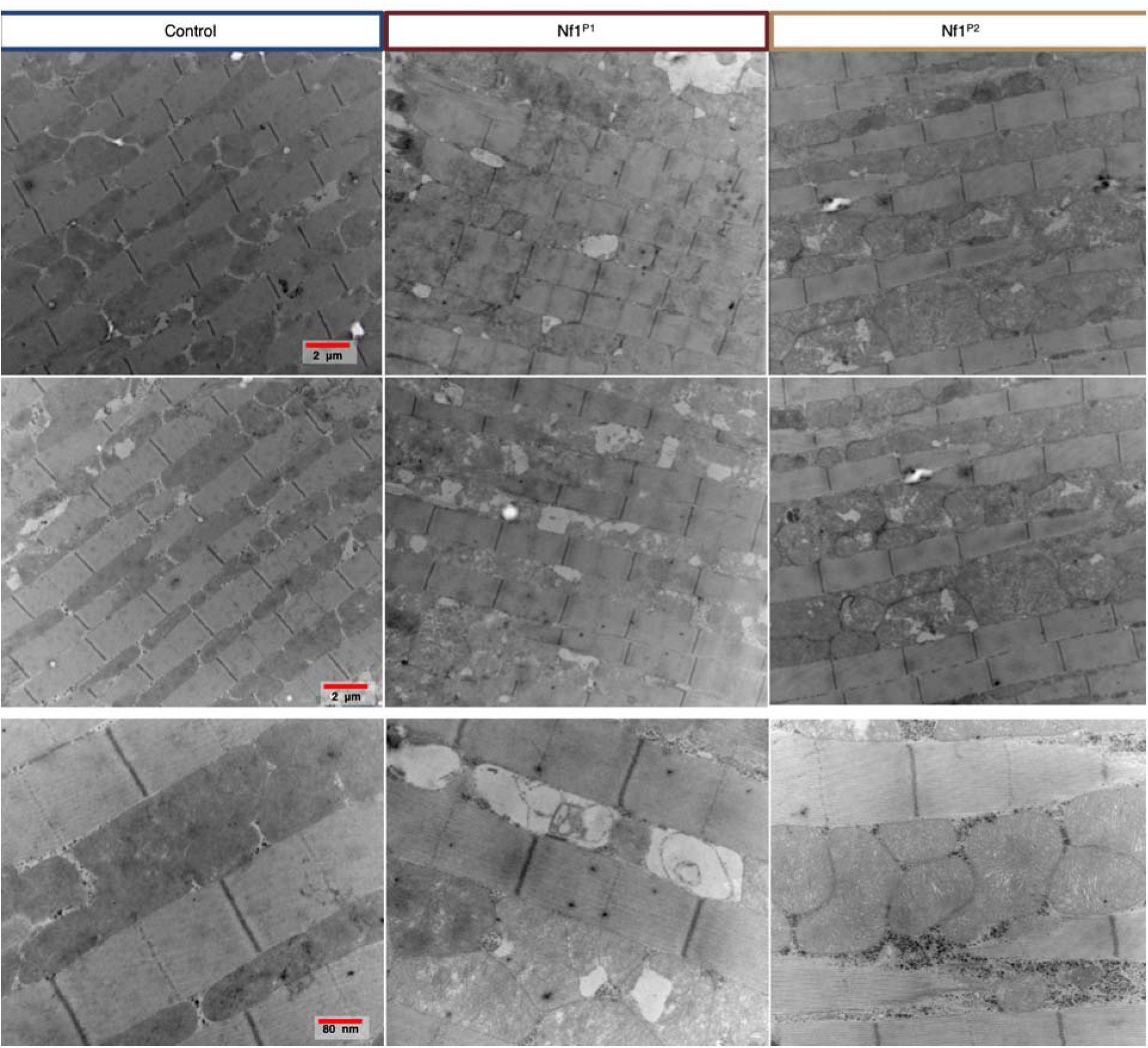
Electron microscopy of *Nf1*-null mutants reveals mitochondrial abnormalities. **(a)** Additional examples of mitochondrial abnormalities in *Nf1*-null mutants. Scale bar = 2 um (top rows), 80 nm (bottom row). Fola you should look for a more representative high-res EM of NF1(p2). The image you show shows highly pack normal mitochondrial with lots of glycogen (black dots) around.

**Extended Data Figure 6.**
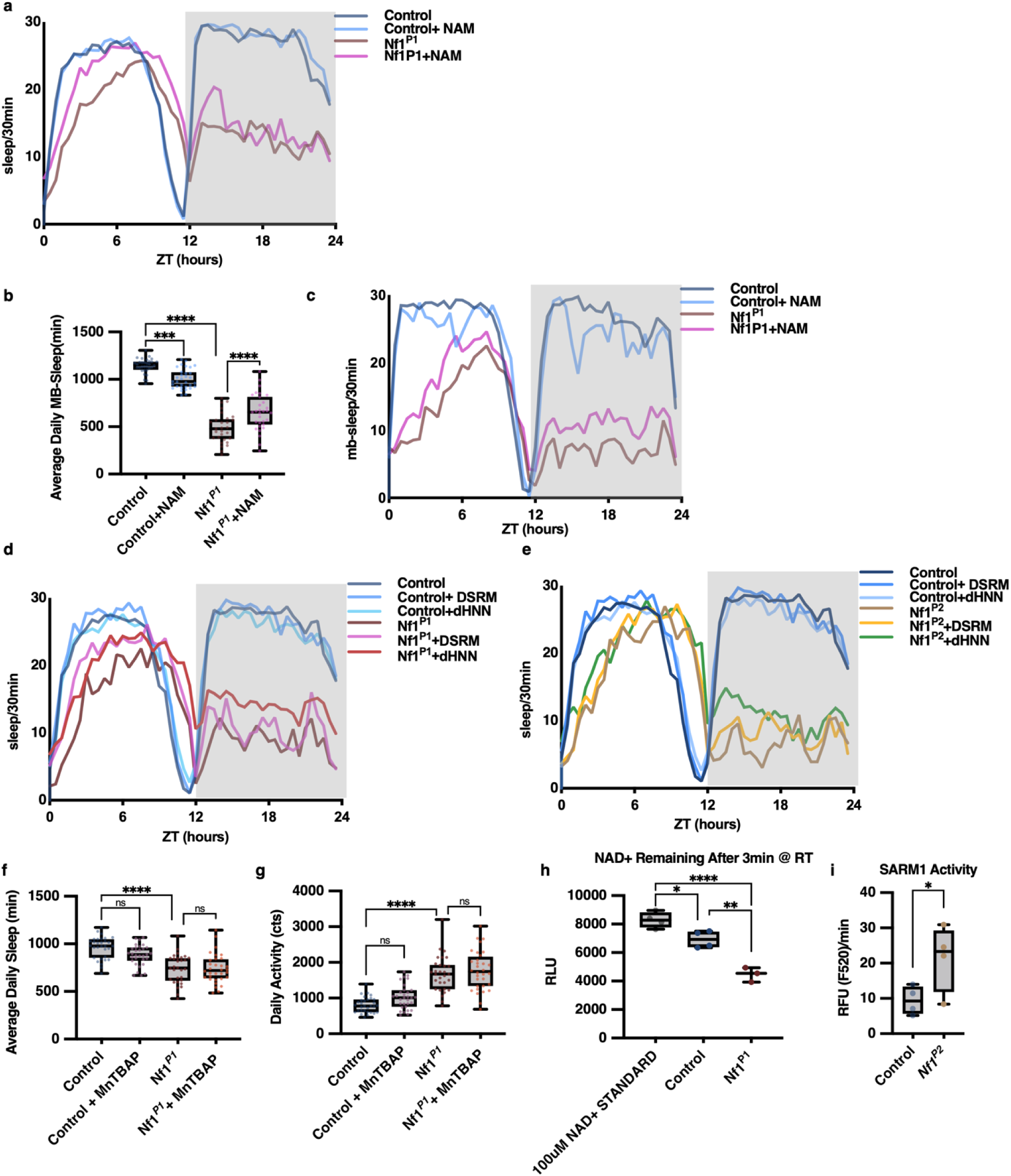
Rescue of Nf1 sleep phenotypes by targeting NAD and SARM1 metabolism. (a) Average sleep traces of *Nf1*-null animals who have been raised on 10mM NAM. **(b)** NAM treatment during development improves sleep duration in *Nf1*-null animals as measured with multibeam infrared actigraphy (n = 32,32,29,31 [L to R], F (3, 120) = 147.8 P<0.0001). **(c)** Averaged sleep traces of *Nf1*-null animals receiving 10mM NAM during development using multibeam infrared actigraphy. (d) Averaged sleep traces of *Nf1*^P1^ animals receiving 10 uM dHNN or 50uM DSRM treatment during the assay period. (e) Averaged sleep traces of *Nf1*^P2^ animals receiving 10 uM dHNN or 50uM DSRM treatment during the assay period. **( (f)** Treatment with the SOD mimetic MnTBAP does not improve sleep duration in *Nf1*-null animals (n = 31,31,30,32 [L to R], F (3, 120) = 39.54 P<0.0001). **(g)** Treatment with the SOD mimetic MnTBAP does not improve sleep duration in *Nf1*-null animals (n = 31,31,30,32 [L to R], F (3, 120) = 21.10 P<0.0001). **(h)** Addition of NAD+ to *Nf1*-null lysate results in immediate depletion. Quantification of NAD+ depletion in *Nf1*-null lysate 3 min after [NAD+] spiked to 100uM NAD+ (n = 3-4 technical replicates F (2, 8) = 40.51 P<0.0001). **(i)** SARM1 activity is increased in Nf1*^P2^* animals as measured by the SARM1-dependent enzymatic conversion of PC6 to PAD6(n= 1 biological replicate, two-tailed t-test t(6) = 2.338, P = 0.0290)

